# TANGO1 and SEC12 are co-packaged with procollagen I to facilitate the generation of large COPII carriers

**DOI:** 10.1101/406348

**Authors:** Lin Yuan, Samuel J Kenny, Juliet Hemmati, Ke Xu, Randy Schekman

## Abstract

Large COPII-coated vesicles serve to convey the large cargo procollagen I (PC1) from the endoplasmic reticulum (ER). The link between large cargo in the lumen of the ER and modulation of the COPII machinery remains unresolved. TANGO1 is required for procollagen (PC) secretion and interacts with PC and COPII on opposite sides of the ER membrane, but evidence suggests that TANGO1 is retained in the ER, and not included in normal size (<100nm) COPII vesicles. Here we show that TANGO1 is exported out of the ER in large COPII-coated PC1 carriers, and retrieved back to the ER by the retrograde coat, COPI, mediated by the C-terminal RDEL retrieval sequence of HSP47. TANGO1 is known to target the COPII initiation factor SEC12 to ER exit sites through an interacting protein, cTAGE5. SEC12 is important for the growth of COPII vesicles, but it is not sorted into small budded vesicles. We found both cTAGE5 and SEC12 were exported with TANGO1 in large COPII carriers. In contrast to its exclusion from small transport vesicles, SEC12 was particularly enriched around ER membranes and large COPII carriers that contained PC1. We constructed a split GFP system to recapitulate the targeting of SEC12 to PC1 via the luminal domain of TANGO1. The minimal targeting system enriched SEC12 around PC1 and generated large PC1 carriers. We conclude that TANGO1, cTAGE5, and SEC12 are co-packed with PC1 into COPII carriers to increase the size of COPII thus ensuring the capture of large cargo.

## Introduction

The COPII protein complex is required for the ER export of most secretory proteins and transmembrane proteins destined for the plasma membrane (1). COPII subunits cooperate to generate transport vesicles that carry secretory cargos to the Golgi apparatus (2). The ER membrane protein SEC12 recruits and activates the small GTPase SAR1 by catalyzing a GDP to GTP exchange (3). SAR1-GTP extends an N-terminal amphipathic helix that embeds in the ER membrane to initiate a vesicle bud (4). SAR1-GTP also recruits the inner layer of COPII coat proteins SEC23/24, which in turn recruit the outer layer COPII coat proteins SEC13/31. Assembled COPII envelops a membrane bud, and on vesicle fission, the coat is shed by the acceleration of GTP hydrolysis by SAR1 (5).

COPII-coated vesicles are usually observed as small vesicles with diameters under 100nm, of sufficient size to transport most secretory cargos (2, 6, 7). Some secretory cargos, such as the 300nm long procollagen I (PC1) rigid rod, require COPII for their secretion, but are seemingly too large to be accommodated by conventional COPII-coated vesicles (8–10). Recently, we reported the existence of large COPII-coated PC1 carriers with diameters above 300nm visualized using correlated light electron microscopy (CLEM), stochastic optical reconstruction microscopy (STORM), and live cell imaging in multiple PC1-secreting cultured human cell lines (11). Although the formation of large ER transport vesicles is promoted by monoubiquitylation of the large subunit of the outer coat, SEC31A (12), the molecular link between ubiquitylation and the change in COPII polymerization is not understood.

Secretory proteins are collected into nascent COPII buds through the intervention of a membrane sorting receptor (13). One such receptor, TANGO1 (MIA3), is required for procollagen (PC) secretion through its interaction with PC1 in the ER lumen and the SEC23/24 subunits on the cytoplasmic surface of the ER (14, 15). The luminal SH3 domain of TANGO1 interacts with the PC-specific chaperone HSP47, which accompanies folded PCs to the cis-Golgi or the ER-Golgi intermediate compartment (ERGIC) in large COPII carriers (Fig. 1 A) (11, 14, 16). TANGO1 also forms a stable complex with two other transmembrane proteins: cTAGE5 and the COPII initiating factor, SEC12 (17–19) (Fig. 1 A). The cytosolic proline-rich domains (PRD) of both TANGO1 and cTAGE5 also interact directly with the COPII inner coat protein, SEC23 (14, 20) (Fig. 1 A). Therefore, TANGO1 has the molecular features of a COPII receptor for large cargo.

**Figure 1.**
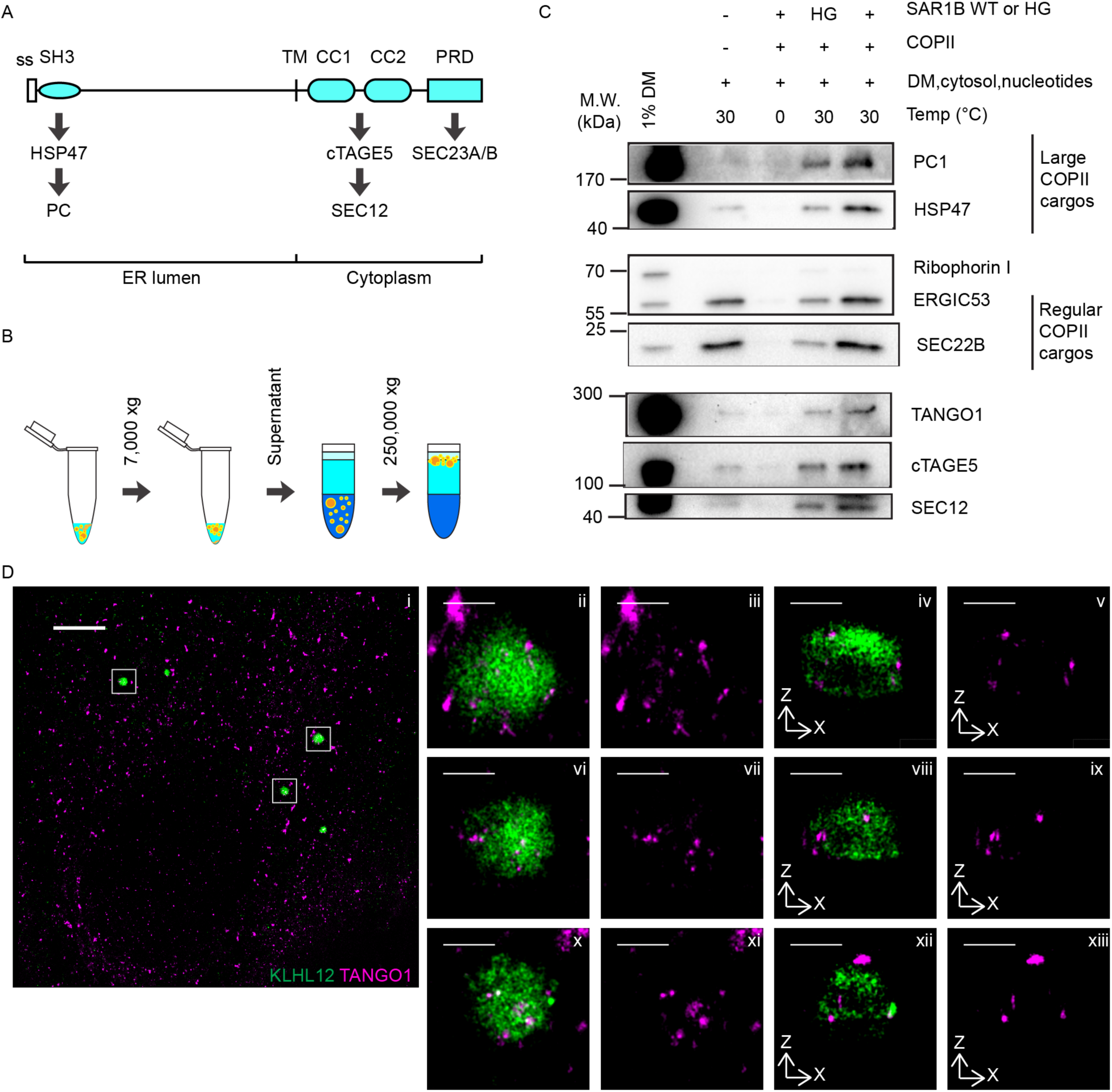
TANGO1 is co-packaged with PC1 into large COPII carriers. (A) Schematic representation of TANGO1’s domain structure with information on relevant interactions. The N-terminal SH3 domain interacts with the PC-specific chaperone HSP47 which binds to folded PCs (16). The coiled-coil (CC) domains of TANGO1 form a stable complex with cTAGE5 and SEC12 (17–19). The proline-rich domain (PRD) of TANGO1 interacts with the COPII inner coat protein SEC23 (14, 20). (B) Scheme depicting the isolation of COPII carriers from cell-free budding reactions as previously described (7,17). Briefly, COPII vesicles were generated by incubating a reaction containing donor membrane prepared from IMR-90 cells, purified recombinant COPII proteins (1µg of SAR1B wt or H79G, 1µg of SEC23A/24D, 1µg of SEC13/31A), nucleotides, and 2µg/µl HT1080 cytosol at 30°C for 1h. Vesicles in 7000xg supernatant fractions from budding reactions were isolated by flotation. (C) TANGO1, cTAGE5, and SEC12 in COPII carriers were detected by immunoblotting aliquots of the top float fractions of reactions conducted under different conditions. Donor membrane (DM) was included as an input control. PC1 and HSP47 are captured into large COPII-coated PC1 carriers and serve as positive controls for large COPII carriers. ERGIC53 and SEC22B are found in conventional COPII vesicles and serve as further controls for COPII vesicles. Ribophorin I is an ER resident protein that serves as a negative control. (D) Dual color 3D-STORM images of a KI6 cell containing multiple large COPII coated vesicles. Overview of the cell is shown in (i), and inserts are enlarged and shown in (ii-xiii). TANGO1 (magenta) co-localized with KLHL12-FLAG (green) which coats large COPII vesicles in XY maximum projections (ii,iii,vi,vii,x,xi) or XZ cross-sections (iv,v,viii,ix,xii,xiii). Scale bars: 5µm in D i; 500nm in D ii-xiii.

Cargo receptors, such as ERGIC53 (LMAN1 or p58), are efficiently sorted into COPII vesicles for anterograde trafficking and are then recycled back to the ER in COPI vesicles (13). In a living cell, newly-generated COPII vesicles are efficiently targeted for fusion with their destination organelles, making it challenging to isolate and characterize vesicle cargo proteins. Fortunately, COPII-coated vesicles can be generated from purified components in a cell-free reaction and thus are more easily isolated; this has proven to be a powerful means to detect the incorporation of cargo receptors, such as ERGIC53 and Erv29p (6, 21). Although this reaction typically generates small COPII vesicles, we recently developed an alternative budding and isolation protocol in order to detect the capture of PC1 into large COPII-coated carriers which were evaluated by immunoblotting, structured illumination microscopy (SIM), flow cytometry, and thin-section transmission electron microscopy (TEM) (11, 22).

Here, we examine large COPII-coated PC1 carriers generated in a cell-free reaction and observe that TANGO1, cTAGE5, and SEC12 are co-packed with PC1. TANGO1 and SEC12 were also observed on endogenous large COPII-coated PC1 carriers by 3D-STORM super-resolution microscopy. Exported TANGO1 is recycled back to the ER with HSP47, a process dependent on the COPI coat, consistent with the typical itinerary of a COPII cargo sorting receptor. In contrast to the exclusion of SEC12 from regular COPII vesicles, SEC12 was enriched around PC1 in budding membrane profiles. To test the effect of actively targeting SEC12 to large cargo by TANGO1, we reconstituted the targeting with minimal components in cultured cells and observed the formation of large PC1 carriers containing SEC12. Thus, our data reveal a novel mechanism in which the large cargo receptor, TANGO1, coordinates the formation of large COPII carriers by actively targeting SEC12 to ER exit sites engaged in the capture of large cargo.

## Results

### TANGO1, cTAGE5, and SEC12 are co-packaged into large COPII carriers along with PC1

The large transmembrane protein TANGO1 is poised to be a COPII receptor for large PC cargo as it interacts with COPII and PC on opposite sides of the ER membrane (Fig. 1 A)(14, 16, 18–20). To test whether TANGO1 is incorporated into COPII carriers with large cargos, we devised a cell-free reaction to generate large COPII-coated PC1 carriers from purified components (11, 22). Following the completion of the reaction, an alternative purification method was used, where donor membrane was sedimented in a 10min centrifugation at 7000xg, and vesicles in the supernatant were separated from soluble components by buoyant density flotation in a step gradient (Fig. 1 B).

Using this isolation method, we previously demonstrated that the capture of PC1 into large COPII-coated membrane carriers was dependent on the presence of COPII coat proteins as well as GTP hydrolysis by the COPII subunit SAR1 (11). Consistent with our previous report, the large cargo PC1, the collagen-specific chaperone HSP47, and the control COPII cargos ERGIC53 and SEC22B were observed in the floated fraction produced in a reaction containing membranes, COPII and nucleotide (Fig. 1 C). Cargo capture was dependent on COPII, and it was reduced in an incubation containing a GTPase mutant, SAR1 H79G (Fig. 1 C). Thus, the COPII-dependent generation of PC1 carriers was recapitulated in the cell-free assay.

Under optimal conditions of temperature and recombinant COPII proteins, all three components of the TANGO1/cTAGE5/SEC12 complex were detected by immunoblotting in the floated fraction (19) (Fig. 1 A, C). The amount of TANGO1, cTAGE5, and SEC12 detected in the floated fraction decreased when recombinant COPII was omitted or when the SAR1B H79G mutant was used in place of wild-type SAR1B. Notably, the export of PC1 and proteins implicated in PC1 secretion, namely HSP47, TANGO1, cTAGE5, and SEC12, showed a higher dependency on recombinant COPII compared to cargos of conventional small COPII vesicles, as more dramatic decreases were observed in PC1, HSP47, TANGO1, cTAGE5 and SEC12 when recombinant COPII was omitted from the reaction (Fig. 1 C, compare lanes 2 from the left to the rightmost lanes). Thus, the export of TANGO1, cTAGE5 and SEC12 share the requirements for sorting into large COPII carriers along with PC1 and standard cargo.

Previous work has shown that mild overexpression of KLHL12, a substrate adaptor of the cullin 3 (CUL3) ubiquitin ligase, leads to the formation of large COPII vesicles and an enhanced rate of traffic of PC1 from the ER to the Golgi complex (11, 12). We engineered human cells (KI6) that overexpress KLHL12 under doxycycline-controlled transcriptional activation and found 7.5h of induction was optimal to observe large PC1-containing COPII structures (11). These large COPII structures were resolved by 3D-STORM revealing hollow spheres of KLHL12 and the COPII coat protein SEC31A that encapsulated the large cargo PC1 (11). We also visualized endogenous large COPII-coated PC1 carriers in Saos-2 cells using 3D-STORM and showed SEC31A and endogenous KLHL12 enveloping endogenous PC1 (11). KI6 and Saos-2 cells were used interchangeably in this study, and both SEC31A and KLHL12 were used as markers for large COPII carriers. To test whether TANGO1 was incorporated into large COPII carriers in cells, we performed immunofluorescence labeling of TANGO1 and KLHL12 in KI6 cells induced for 7.5h as before. Large, hollow spheres of KLHL12 >300nm in diameter were observed, as previously reported (11), with TANGO1 contained within the large COPII carriers (Fig. 1 D). The super-resolution visualization and cell-free biochemical results confirm each other.

### TANGO1 is recycled with HSP47 by a COPI-dependent process

Similar to other cargo receptors like ERGIC53, TANGO1 localizes at ER exit sites (ERES) in cells at steady state. Unlike most cargo adaptors, TANGO1 does not contain a C-terminal KKXX or KDEL retrieval signal. Alternatively, the collagen chaperone HSP47 may be responsible for retrieval of this receptor as it interacts with the C-terminal SH3 domain of TANGO1 (14). HSP47 recognizes the folded triple-helical domain of PCs in the ER (23), and serves as a chaperone to convey cargo in large COPII carriers to the ERGIC or cis-Golgi compartment (11). Subsequently, cargo is released due to the lower pH in the lumen of ERGIC or cis-Golgi, and HSP47 is recycled back to the ER via its C-terminal RDEL sequence (24, 25). Efficient recycling results in the steady-state localization of HSP47 to the ER. When the C-terminal RDEL sequence is deleted, HSP47ΔRDEL is readily secreted (24). To test whether TANGO1 is retrieved via its interaction with HSP47, we overexpressed a StrepII tagged HSP47ΔRDEL and observed that in cells that had secreted this fusion, TANGO1 either mislocalized to the Golgi apparatus (Fig. 2Bii) or was dispersed more broadly (Fig. 2A, Bi). These results showed that the interaction between TANGO1 and HSP47 influences the retrieval of TANGO1.

**Figure 2.**
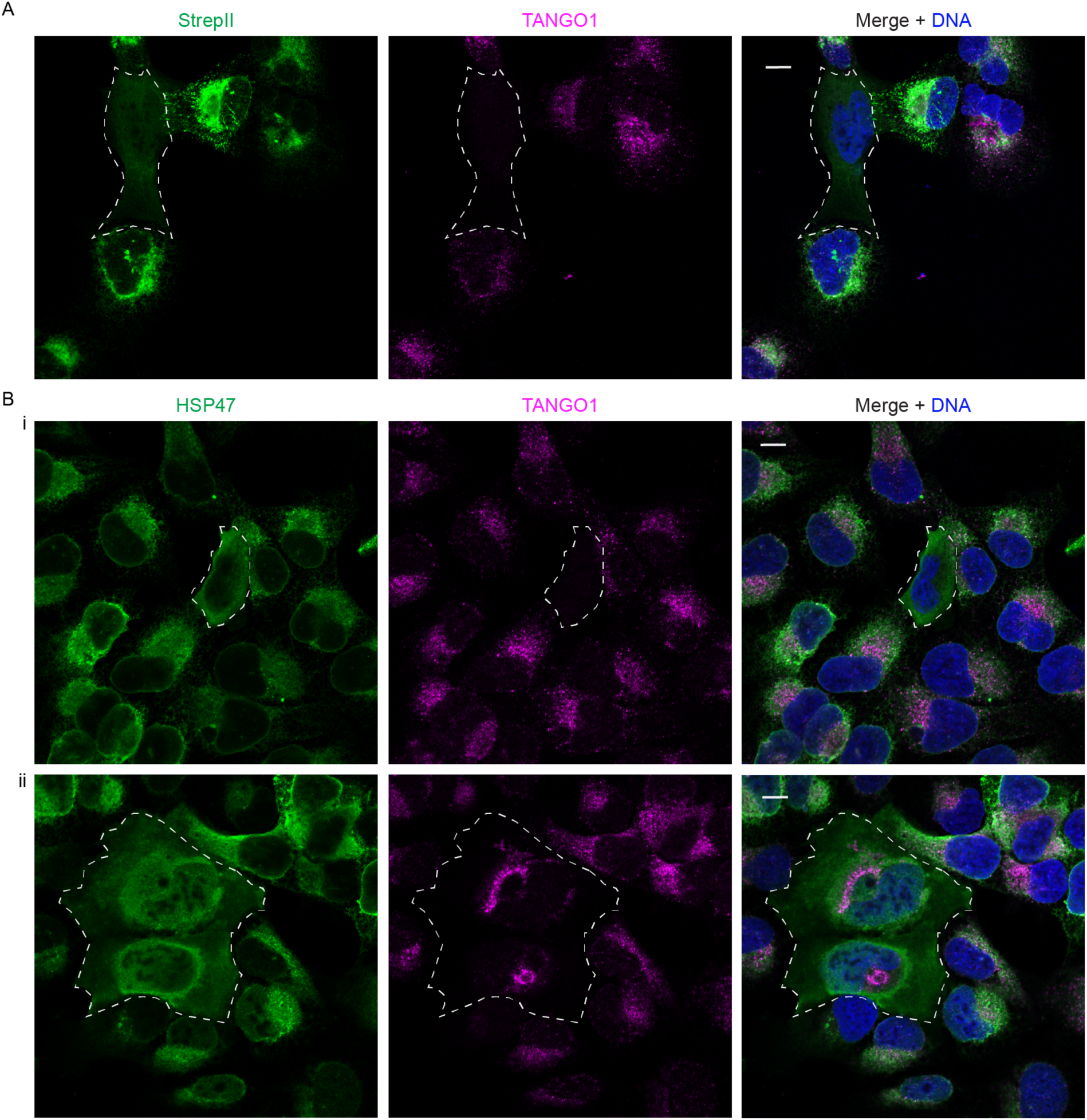
TANGO1 is retrieved with HSP47 via its c-terminal RDEL motif. Confocal microscopy images of U-2OS cells transfected with StrepII-HSP47ΔRDEL and labeled with a fluorescent antibody against the StrepII tag (A; green) or HSP47 (B; green) and TANGO1 (magenta). In wildtype cells, TANGO1 localized at ERES and HSP47 localized in the ER. TANGO1 mis-localization is observed following the overexpression and secretion of StrepII-HSP47ΔRDEL in cells marked inside dotted lines. In these cells, HSP47 (green) no longer showed ER localization, but rather appeared to localize at the cell surface; TANGO1 (magenta) no longer localized to the ERES, but in some cells localized to the Golgi membrane (2Bii). Scale bars: 10µm.

To test whether TANGO1 is recycled back to the ER by COPI, we depleted COPI in cells with siRNA that targets coatomer subunit δ (*ARCN1* gene) (26). In cells that were depleted of COPI, we observed accumulation of TANGO1 around concentrated HSP47 structures (Fig. 3 A). This localization was not observed in cells transfected with negative control siRNA, showing that this phenotype was due to COPI depletion (Fig. 3 A). Alternatively, COPI trafficking is blocked by overexpressing a GTP-locked ARF1 Q71L mutant (26). In cells that expressed ARF1 Q71L-GFP, TANGO1 accumulated around HSP47 puncta similar to the phenotype observed with siRNA knockdown of coatomer subunit δ (Fig. 3 B). Together, these data suggest that the retrieval of TANGO1 depends on ARF1-GTP hydrolysis and COPI budding.

**Figure 3.**
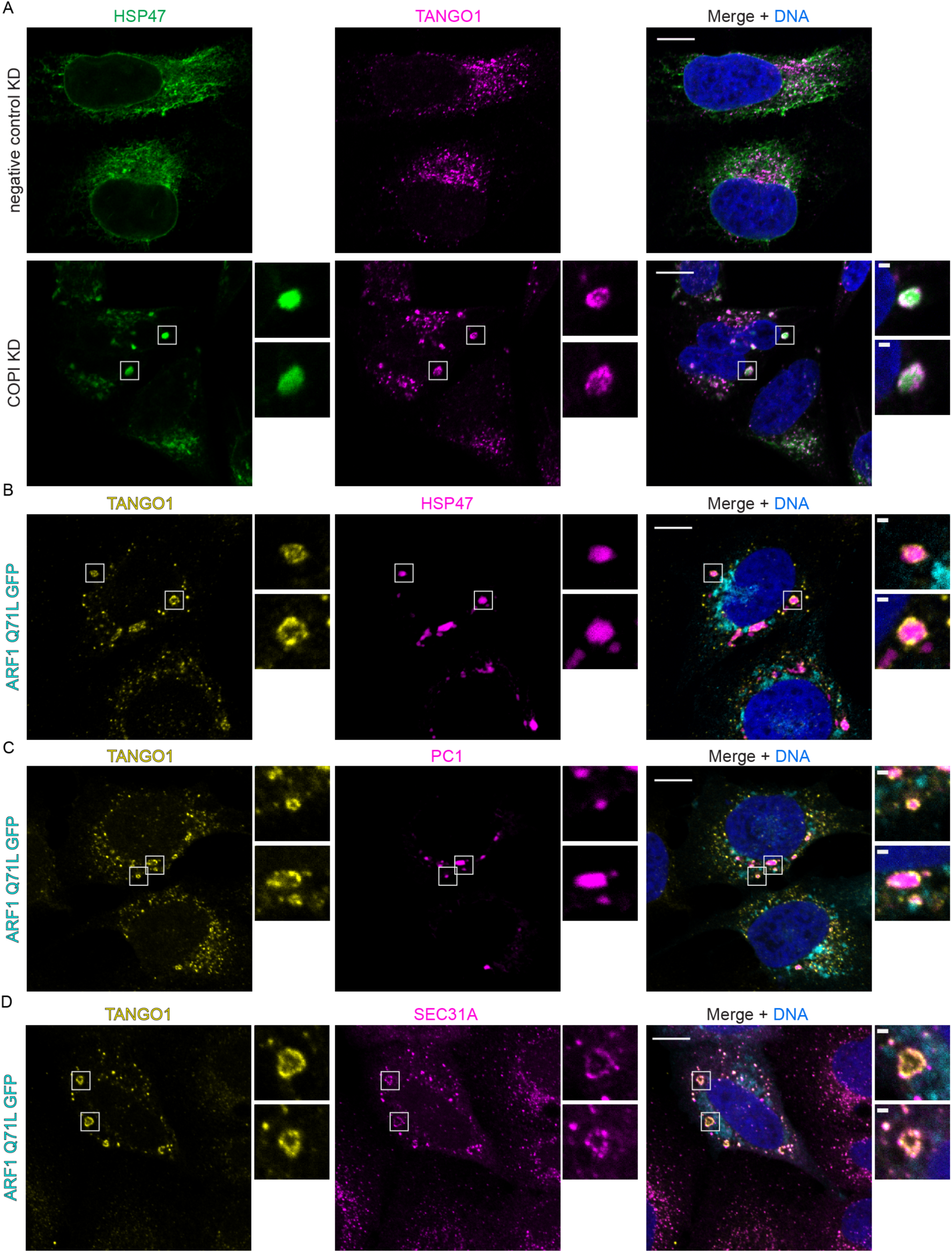
TANGO1 localizes around large COPII carriers in cells depleted of COPI. (A) Confocal images of U-2os-wt-c11 cells transfected with negative control siRNA or siRNA that targeted coatomer subunit δ (ARCN1 gene) for 48h followed by immunofluorescence labeling of TANGO1 (magenta) and HSP47 (green). TANGO1 was observed around large HSP47 puncta in cells depleted of COPI. Magnified inserts show two examples of such structures. (B-D) Confocal images of U-2os-wt-c11 cells expressing ARF1 Q71L GFP (cyan) were labeled by immunofluorescence targeting of TANGO1 (yellow) and HSP47 (magenta) in (B) or PC1 (magenta) in (C) or SEC31A (magenta) in (D). Scale bars: 10µm in overviews and 1µm in magnified inserts.

We further characterized the compartment where TANGO1 accumulated in cells depleted of COPI. TANGO1-decorated HSP47 puncta appeared not to co-localize with ARF1 Q71L-GFP, which localizes in the ERGIC (26). Instead, TANGO1 accumulated around PC1 puncta and colocalized with the COPII outer coat protein SEC31A (Fig. 3 C,D). These exceptionally large COPII structures were much bigger than the functional carriers we observed by STORM and CLEM (11), and were readily resolved by confocal microscopy.

### SEC12 is enriched in large COPII-coated PC1 carriers

We were particularly intrigued by the detection of SEC12 in COPII carriers in our cell-free reaction (Fig. 1 C), because this protein is not normally sorted into small COPII vesicles (2, 3). A recent study reconstituted the cytosolic and transmembrane domains of the yeast Sec12p and the transmembrane COPII cargo Bet1p on a thick planar lipid bilayer that allowed collection of cargo molecules into curved membrane buds but did not support vesicle scission (27). When COPII proteins (Sar1p, Sec23p/24p, Sec13p/31p) and GTP were supplemented to the planar lipid bilayer containing Sec12p and Bet1p, COPII coat proteins (Sec23/24, Sec13/31) polymerized into clusters with the cargo Bet1p, resembling pre-budding complexes at the ERES (27). Consistent with our own earlier results using native ER membranes as a template for vesicle budding (2), Sec12p was excluded from the reconstituted COPII-cargo clusters, suggesting that it is intrinsically excluded from regular COPII pre-budding complexes and thus regular COPII vesicles (27).

We hypothesize that the lateral organization of SEC12 may be controlled to allow the recruitment of SAR1-GTP onto large COPII carriers where it may serve to sustain the polymerization of the coat onto an enlarged surface (2). To test this hypothesis, we devised a fractionation scheme to separate large and regular COPII carriers generated by the cell-free reaction (Fig. 4 A). After the incubation was completed, cell-free reactions were centrifuged for 10min at 7000xg to sediment donor membranes. The supernatant fraction was taken and further sedimented through a step Optiprep density gradient at 250,000xg for 1h to separate COPII carriers of regular and large cargo. Fractions taken after sedimentation were used as input for flotation, which would separate the membrane from soluble components, and the floated sedimentation fractions were analyzed by immunoblot (Fig. 4 B). Most PC1 sedimented to the interphase between 0% and 7.5% Optiprep in fraction 2, a relatively low buoyant density position in relation to typical COPII vesicles. In contrast, most regular COPII markers ERGIC53 and SEC22B sedimented to the interphase between 7.5% and 18% Optiprep, fraction 4, a more typical high buoyant density position (11). Thus, the physical properties of PC1-containing and regular cargo COPII vesicles appear to differ.

**Figure 4.**
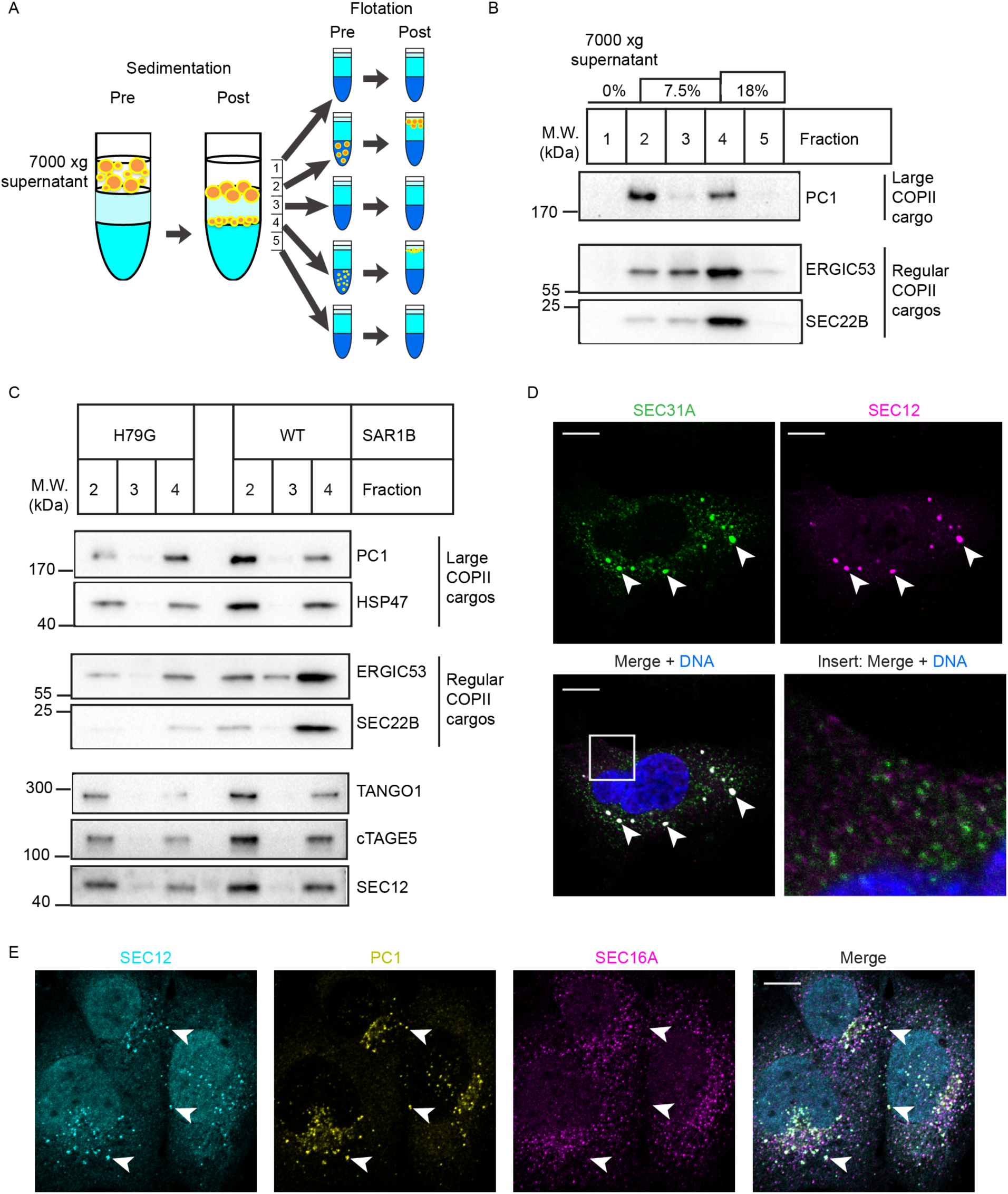
SEC12 is enriched in large COPII-coated PC1 carriers. (A) Schematic representation of the fractionation procedure used to separate small and large COPII carriers. Supernatant after 7000xg centrifugation from a vesicle-budding reaction was overlaid onto a step gradient consisting of 7.5% and 18% OptiPrep. The gradient was centrifuged at 250,000xg for 1h to separate large from regular cargo-containing COPII carriers. Fractions (numbered 1-5) taken after sedimentation were used as inputs for flotation to separate vesicles from soluble contents. (B) Analysis of the large cargo PC1 and regular COPII cargos ERGIC53 and SEC22B in buoyant membrane collected from sedimentation fractions post-flotation. (C) Budding reactions were supplemented with wild-type (WT) or H79G mutant SAR1B. Buoyant membrane from relevant sedimentation fractions were immunoblotted for TANGO1, cTAGE5, and SEC12. PC1 and HSP47 serve as markers for COPII-coated PC1 carriers. ERGIC53 and SEC22B serve as markers for regular COPII vesicles. (D) Confocal image of KI6 cells that were induced for KLHL12 overexpression for 7.5h and immunolabeled with a fluorescent antibody against SEC31A (green) and SEC12 (magenta). (E) Confocal immunofluorescence images of SEC12 (cyan), PC1 (yellow), and SEC16A (magenta) in Saos-2 cells. Arrows point to examples of large SEC12 puncta that colocalized with PC1 but not SEC16A. Scale bars: 10µm in (D,E).

To test whether TANGO1, cTAGE5, and SEC12 are packaged into COPII-coated PC1 carriers, we used immunoblot to detect these proteins in the relevant buoyant density fractions. Co-fractionation of TANGO1, cTAGE5, and SEC12 with large PC1 carriers was observed, as they were more abundant at the 0% to 7.5% interphase (Fig. 4 C). When the SAR1B H79G mutant was supplemented to inhibit COPII budding, less PC1 and HSP47 were detected at the 0% to 7.5% interphase. The enrichment of TANGO1, cTAGE5, and SEC12 in this fraction was also sensitive to the SAR1B H79G mutant, confirming that TANGO1, cTAGE5, and SEC12 were incorporated into low buoyant density COPII-coated PC1 carriers in a manner dependent upon GTP hydrolysis by SAR1.

Previously, we found that small COPII vesicles were about 10 to 20-fold more prevalent than large COPII-coated PC1 carriers in the 7000xg supernatant fraction as quantified by flow cytometry and nanoparticle tracking analysis (11). Although the immunoblot in Fig. 3C suggested that SEC12 was only somewhat enriched in the low vs. the high buoyant density membranes, the relative enrichment per COPII vesicle may be substantially greater. We examined the localization of SEC12 in KI6 cells after 7.5h of induced overexpression of KLHL12, conditions that produce large COPII-coated PC1 (11). Using confocal microscopy, we observed large SEC12 puncta that co-localized with large SEC31A puncta (Fig. 4 D). Smaller SEC12 puncta, possibly representing ERES for regular cargo, were also observed at lower signal intensity (18) (Fig. 4 D, inset). Small SEC31A puncta represent both ERES and small COPII vesicles. Populations of small SEC31A puncta that did not co-localize with SEC12 were observed, possibly representing free small COPII vesicles that excluded SEC12 (Fig. 4 D, inset). Large SEC12 puncta were also observed by confocal microscopy in PC1-secreting Saos-2 cells not overexpressing KLHL12 (Fig. 4 E). As mammalian SEC12 is known to localize at the ERES, we also included the scaffold protein SEC16A as a marker for ERES. To stimulate ER export of PC1, we treated Saos-2 cells with ascorbate for 30min prior to fixation. Ascorbate is a co-factor for prolyl-hydroxylase which is required for PC trimerization, thus its addition stimulates PC1 secretion. We observed large and densely labeled SEC12 puncta that were predominantly positive for PC1, and many of these large SEC12 puncta did not co-localize with SEC16A suggesting they were free large COPII carriers of PC1 (Fig. 4 E, arrows). Large SEC12 puncta that were positive for both PC1 and SEC16A were also observed, suggesting that SEC12 also localized to PC1-containing ERES.

### SEC12 is localized around PC1 in large COPII structures

We next employed 3D-STORM to resolve the large SEC12 puncta observed by confocal microscopy. Three classes of ultra-structures were revealed when large SEC12 puncta over 300nm in diameter were examined in 3D (Fig. 5 A). The first class of large SEC12 structures were hollow spheres, similar to what we previously observed for coat component SEC31A in large COPII carriers (11) (Fig. 5 A iii). To study the location of PC1 and the COPII coat with respect to SEC12, we performed three-color 3D STORM imaging on large SEC12/PC1/SEC31A puncta (Fig. 5 B iii). PC1 was resolved to be inside of hollow cavities and entirely encapsulated by SEC12 and SEC31A, suggesting that these SEC12 hollow spheres were large COPII-coated PC1 carriers (Fig. 5 C iii). The second class of large SEC12 structures were cup-shaped structures (Fig. 5 A ii), which were also previously reported with SEC31A (11). These structures appeared to be nascent budding events at the ERES, as the cup-shaped SEC12/KLHL12 co-localized structure only partially enveloped PC1 (Fig. 5 B ii; C ii). A third class of large SEC12 structures appeared to be flat discs with little curvature (Fig. 5 A i), which were not observed when the localization of the COPII outer coat protein SEC31A was analyzed by 3D-STORM. Although these SEC12 flat discs co-localized with PC1 without a discernible pattern in maximum XY projections, PC1 localized to only one side of the SEC12 flat discs when the 3D structure was examined (Fig. 5 B i). The SEC12/PC1 flat discs possibly represented PC1-containing ERES before the recruitment and activation of SAR1 (Fig. 5 C i), which would explain why little SEC31A was observed overlapping with SEC12, given that the SEC13/31 outer coat is recruited after the activation of SAR1 and the recruitment of the inner coat (28). These 3D-STORM data supported our biochemical analyses of large COPII-coated PC1 carriers generated in a cell-free reaction.

**Figure 5.**
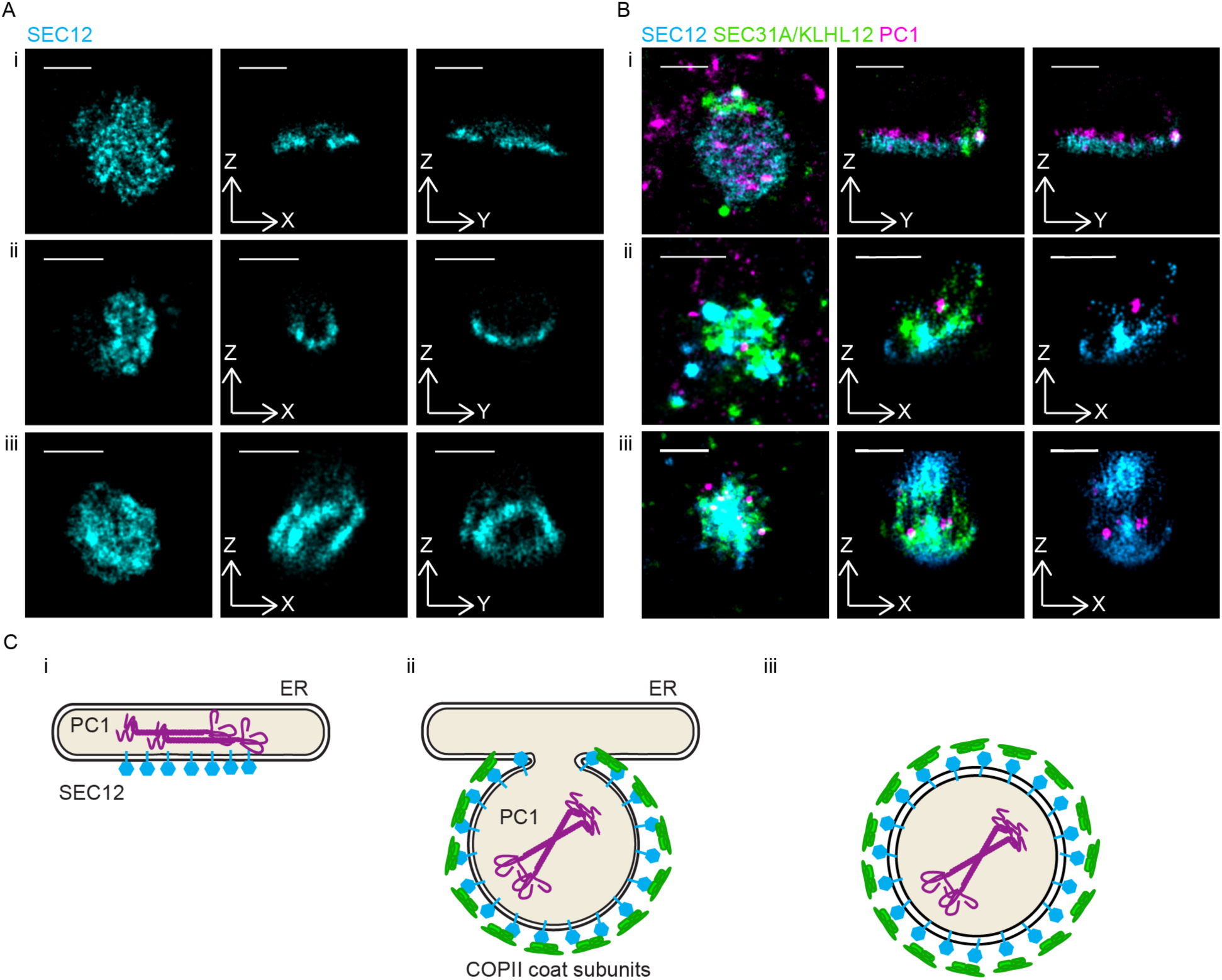
SEC12 is localized around PC1 throughout large COPII vesicle formation. (A) 3D STORM images of single color large SEC12 (cyan) puncta collected from Saos-2 cells. Three representative examples from three classes of ultra-structures (i-iii) are shown in magnified maximum XY-projection (left), virtual cross-sections in XZ (middle), and YZ (right). (B) Three color 3D STORM images of large COPII structures in KI6 cells: SEC12 (cyan), PC1 (magenta), and COPII coat subunit SEC31A (green in Bi, Biii) or KLHL12 (green in Bii). Representative examples of three classes of ultra-structures (i-iii) are shown in three-color merged maximum XY projection (left), three-color merged virtual cross-sections (middle), and two-color merged virtual cross-sections of SEC12 (cyan) and PC1 (magenta) (right). (C) Schematic illustrations of three classes of ultra-structures of SEC12 arranged in putative time progression: (i) enrichment of SEC12 (cyan) around PC1 (magenta) containing ER; (ii) nascent large COPII (green) budding event where SEC12 localizes around membrane-containing PC1; (iii) free large COPII-coated PC1 carrier with enriched SEC12. Scale bars: 500nm.

We deduced a putative temporal progression of large COPII-coated PC1 carrier formation based on the three classes of SEC12 ultra-structures revealed by 3D-STORM (Fig. 5 C). In this speculative timeline, the concentrated targeting of SEC12 to PC1-containing ERES precedes the recruitment of the SAR1 GTPase, which is activated when SEC12 catalyzes its guanine nucleotide exchange (29–31). Because binding of SAR1 to GTP exposes an amphipathic helix that is sufficient to induce membrane curvature and recruit downstream COPII coat subunits to complete vesicle budding, the flat discs of SEC12 may be formed prior to the initiation of curvature (4, 32).

### Active targeting of SEC12 to large cargo increases COPII size

To test whether the active sorting of SEC12 could control the size of COPII carriers, we recapitulated the targeting of SEC12 to PC1-containing ER membrane in cultured cells. Previous reports showed that TANGO1 and cTAGE5 mediate the targeting of SEC12 to ERES, as knocking down either TANGO1 or cTAGE5 resulted in diffuse ER localization of SEC12 (18, 19, 33). We considered the possibility that SEC12 is targeted to PC1-containing ERES mediated by the luminal SH3 domain of TANGO1. This SH3 domain is the only element in the TANGO1/cTAGE5/SEC12 complex that is known to bind PC1 through its interaction with HSP47 (Fig. 1A; Fig. 6 A). To test this hypothesis, we employed a split-GFP targeting and detection scheme based on the self-assembly of complementary fragments of GFP fused to each target protein (29). This method has been shown to drive interaction and to visually localize targets in close proximity. Briefly, GFP11 (the 11th β-strand of GFP) was fused to the C-terminus of 3xFLAG-SEC12 so that it would be exposed on the short ER luminal tail of SEC12; GFP1-10 (the rest of GFP without the 11th β-strand) was fused to the C-terminus of the luminal SH3, and unstructured domains of TANGO1 (TANGO1-lumi) and an HA tag was used as linker between TANGO1-lumi and GFP1-10 (Fig. 6A). When transfected alone, neither construct produced GFP fluorescence, and both showed ER localization, as expected (Fig. 6 B). In cells transfected with both 3xFLAG-SEC12-GFP11 (referred to as SEC12-GFP11) and TANGO1-lumi-HA-GFP1-10 (referred to as TANGO1-lumi-GFP1-10), SEC12-GFP11 was recruited to TANGO1-lumi-GFP1-10 by GFP complementation (Fig. 6 B). To test whether SEC12-GFP11 was targeted to PC1 when complemented with TANGO1-lumi-GFP1-10, we labeled endogenous PC1 by immunofluorescence and observed co-localization of PC1 with the complemented GFP (Fig. 6 C). Complemented GFP signals were often ER-localized and large GFP puncta were found in a subpopulation of GFP positive cells (Fig. 6 B-C, arrows).

**Figure 6.**
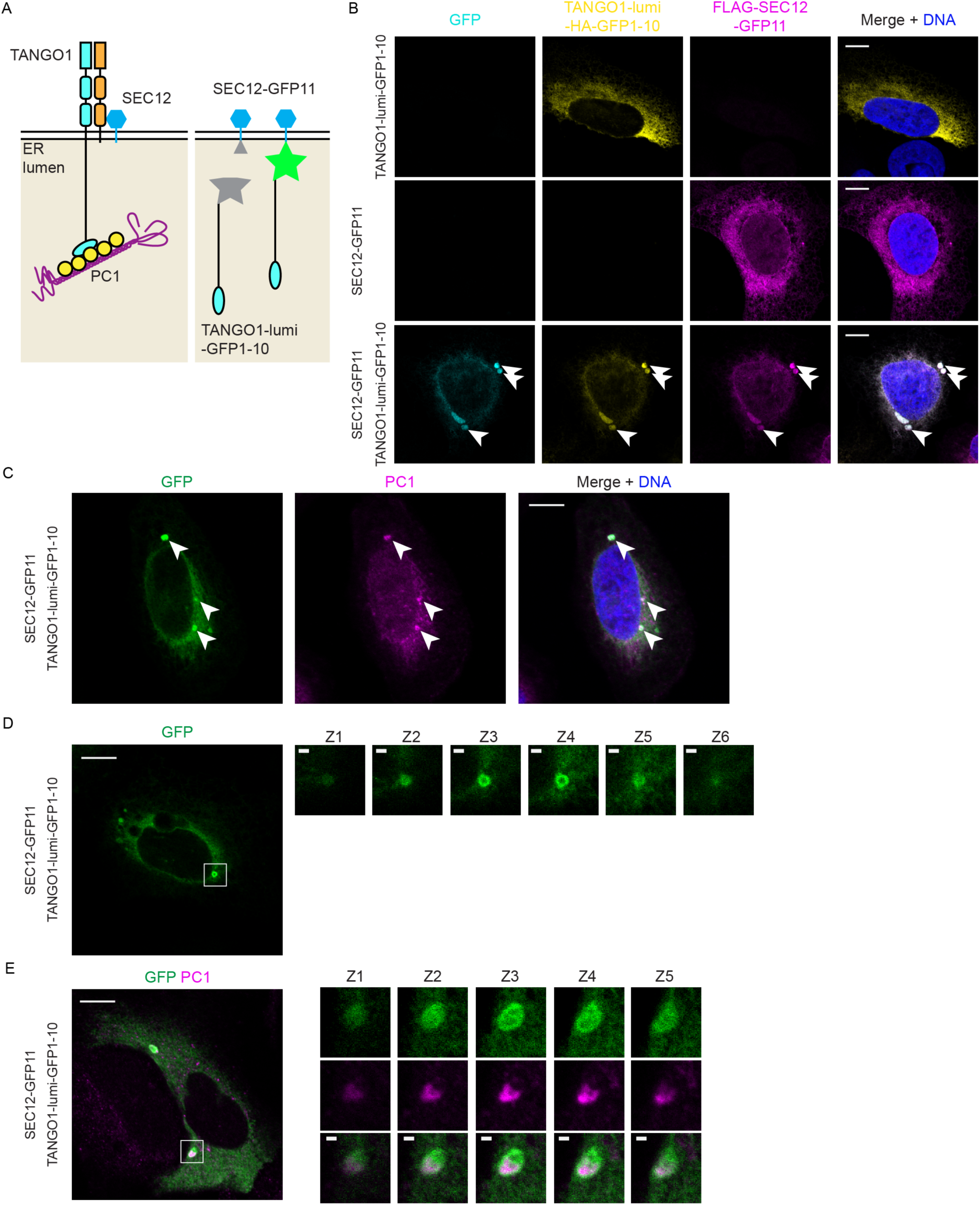
Active targeting of SEC12 to large cargo increases COPII size. (A) Drawings that represent the working model of SEC12 enrichment (left) and the design of split GFP constructs (right). In our working model, the luminal SH3 domain of TANGO1 targets SEC12 to PC1. To recapitulate sorting of SEC12 to PC1, we constructed 3xFLAG-SEC12-GFP11 and TANGO1-lumi-HA-GFP1-10 fusion constructs, where TANGO1-lumi contains TANGO1’s cargo-sensing SH3 domain. (B) GFP complementation brought SEC12 and TANGO1-lumi together. Confocal images of U-2OS-wt-c11 cells that were transfected with TANGO1-lumi-HA-GFP1-10 alone (upper row), 3xFLAG-SEC12-GFP11 alone (middle row), or both (bottom row), complemented GFP (cyan), TANGO1-lumi-HA-GFP1-10 (yellow, IF against HA), and 3xFLAG-SEC12-GFP11 (magenta, IF against FLAG). (C-E) Confocal images of U-2OS-wt-c11 cells transfected with both TANGO1-lumi-HA-GFP1-10 and 3xFLAG-SEC12-GFP11 Cells were immunofluorescently labeled against PC1 (magenta) in (C,E) and imaged in combination with complemented GFP (green). (D,E) Cells were cultured in dialyzed medium, which contained minimal ascorbate. Ascorbate was added to cells for 30 min to stimulate PC1 export. Confocal Z-stacks of magnified insets show hollow spherical complement GFP (green) structures that entirely enveloped PC1 (magenta). Confocal Z stack was taken with a step size of 0.38µm. Scale bars: 10µm (B-E), 1µm (magnified insets in D,E).

We used ascorbate treatment to synchronize the traffic of procollagen in these cells. As regular culture medium contains a trace amount of ascorbate, we cultured cells in dialyzed medium with a minimum level of ascorbate prior to transient transfection with the split GFP constructs. After 20-30min of ascorbate treatment, large GFP structures were observed, many large enough to be visualized as hollow spheres by confocal microscopy (Fig. 6 D). GFP spheres also encapsulated endogenous PC1 as detected by immunofluorescence (Fig. 6 E). The split-GFP-induced large COPII carriers were sometimes large enough to be resolved by confocal microscopy (Z stack of magnified insets, Fig. 6 D,E) and appeared much larger than most endogenous carriers (Fig. 4E). Thus, ectopically targeting overexpressed SEC12 to PC1 further increased the size of large COPII carriers, consistent with our hypothesis that the localization of SEC12 may guide the growth of the coat (3).

We next used an antibody that detects SAR1-GTP to visualize localization to the COPII carriers (34). When wild-type SEC12-GFP11 and TANGO1-lumi-GFP1-10 were transfected together, intense labeling of large SAR1-GTP puncta was observed to co-localize with GFP puncta (Fig. S1). In contrast, SAR1-GTP was not found at GFP puncta when the SEC12 GEF deficient mutant I41A was targeted to PC1 by TANGO1-lumi (3, 35) (Fig. S1). We also observed large SEC31A puncta that co-localized with complemented GFP and PC1, albeit at a low frequency (Fig. S2). These data suggest that active targeting of SEC12 to PC1 can increase the size of COPII vesicles and thus facilitate in the formation of large coated vesicles.

## Discussion

As an essential transport vehicle of the early secretory pathway, COPII vesicles are of strikingly uniform and small size. In search of a mechanism to explain the genetic requirement for COPII to secrete large cargo like PC1 (9, 10), we have previously discovered the existence of large COPII vesicles and established them as *bona fide* carriers of PC1 (11, 12). In this work, we examined their molecular composition and discovered that TANGO1, cTAGE5, and SEC12 are co-packaged with PC1 into COPII carriers (Fig. 1, 4). The ER export of TANGO1 was further supported by the detection of TANGO1 in large COPII-coated carriers in cultured cells as visualized by STORM and the elucidation of a recycling pathway (Fig. 1-3). Further, we provide evidence to suggest that TANGO1 may mediate the sorting of SEC12 to PC1-containing ERES as a novel mechanism to increase the size of COPII carriers (Fig. 4-6).

COPII vesicle formation is a coordinated process centered around the guanine nucleotide status of the small GTPase SAR1, and related to this, several reported mechanisms of COPII size regulation are also linked to the guanine nucleotide status of SAR1 (10, 36). As the first step of COPII formation, SAR1 is recruited to the ER membrane by its GEF SEC12 (29), and the subsequent nucleotide exchange exposes an amphipathic helix which intercalates into the ER membrane, thereby inducing membrane curvature (3, 4). Activated SAR1-GTP recruits the inner coat proteins SEC23/24, where SEC23 is the GTPase activating protein (GAP) for SAR1 (37). The inner coat proteins then recruit the outer coat proteins SEC13/31, where SEC31 acts to stimulate the GAP activity of SEC23 a further 10-fold (5). Regulating the local concentration of SAR1-GTP could change the timing of membrane scission and vesicle size (31). This is supported by the discovery of Sec24p-m11, a mutant that augments Sar1-GTPase hydrolysis and generates smaller-than-normal COPII vesicles (36). Moreover, a mechanistic study of cranio-lenticulo-sutural dysplasia, a disease caused by a deficiency in procollagen export, revealed that the mutation SEC23A M702V inhibits PC secretion by speeding GTP hydrolysis (10)

The large transmembrane protein TANGO1 is essential for the secretion of large cargo including members of the collagen family (14, 15, 38). ER export of TANGO1 was examined previously using a cell-free reaction that supported the generation and detection of small COPII vesicles (14). As TANGO1 was not detected in isolated COPII vesicles, the idea emerged that TANGO1 served to package but not to accompany collagen out of the ER, thus not serving the traditional role of a stoichiometric sorting receptor (14). However, the capture of the large cargo into COPII carriers was not probed in early studies, hence a mechanism of sorting mediated by TANGO1 remained elusive (14). Recently, we reported an improved method that allowed the detection of large COPII-coated PC1 carriers generated in a cell-free reaction (11, 22). In the present study, using this approach, we demonstrated the COPII dependent co-packaging of TANGO1 with PC1 into COPII carriers (Fig. 1 C, Fig. 2 C). This result was confirmed by the detection of TANGO1 on large COPII structures in PC1 secreting cells as visualized by STORM (Fig. 1 D) and the COPI-dependent recycling of TANGO1 (Fig. 3). Consistent with the trafficking defect observed in cells depleted of COPI (Fig. 3), a recent study identified loss-of-function mutations in the *ARCN1* gene which encodes the subunit δ of COPI and causes a human craniofacial syndrome (39). This genetic disease showed similar symptoms to cranio-lenticulo-sutural dysplasia and osteogenesis imperfecta, which are collagen deposition diseases caused by mutations in COPII (9, 39–41).

Similar to TANGO1, the COPII initiating factor SEC12 is not detected in small COPII vesicles generated in cell-free reactions (3, 30). Although the yeast homolog Sec12p is observed to escape the ER and is returned by Rer1p in COPI vesicles, most Sec12p remains in the ER in an *rer1* null mutant, suggesting that the escaped Sec12p accounts for only a small fraction of the total amount of Sec12p in the cell (42, 43). Recently, a truncated recombinant yeast Sec12p missing the luminal domain but retaining cytosolic and transmembrane domains was reconstituted into planar lipid bilayers with minimal yeast COPII components (Sar1p, Sec23p/24p, and Sec13p/31p), and GTP or GMP-PNP (27). This truncated Sec12p resembles mammalian SEC12 homologs, which have short luminal tails, was excluded from cargo-containing COPII buds independent of Sar1-GTP hydrolysis (27). This sorting is likely the result of kinetic segregation from the tightly packed cargo and coat components in COPII buds, analogous to CD45 exclusion from the immunological synapse between T cells and antigen presenting cells (APC) (44). In contrast, here we demonstrate specific enrichment of SEC12 in large COPII carriers and PC1-containing ERES (Fig. 1 C; Fig. 4, 5). As CD45 on T cells can overcome kinetic exclusion by interacting with a binding partner on the APC (44), the specific enrichment of SEC12 may be achieved by forming a stable complex with cTAGE5 and TANGO1 (19), where TANGO1 targets the complex to PC1 via its luminal SH3 domain (Fig. 6A) (14, 16). We tested this possibility by targeting SEC12 to the SH3 domain of TANGO1 using a split-GFP system and observed enrichment of SEC12 around PC1-containing ER membrane and large COPII-coated PC1 carriers (Fig. 6).

To test whether active sorting of SEC12 can lead to an increase in COPII size, we used the split GFP system to reinforce the concentration of SEC12 around PC1-containing ER membrane and observed large COPII-coated PC1 carriers as visualized by confocal microscopy (Fig. 6). The ability of SEC12 to increase vesicle size is likely a result of its catalysis of nucleotide exchange on SAR1, as high levels of SAR1-GTP were detected around complemented GFP signals at ERES (Fig. S1). The importance of a high concentration of activated SAR1 during large cargo secretion was demonstrated in another recent report, where overexpression of wild-type SAR1 but not the GTP locked H79G mutant rescued secretion of collagen VII in cells where the localization of SEC12 was dispersed as a result of cTAGE5 knockdown (37).

In our split GFP system, we reconstituted the active targeting of exogenously overexpressed SEC12 with the luminal domain of TANGO1 and demonstrated a novel role of TANGO1 during ER export of PC1 (Fig. 6). Under normal circumstances, the endogenous enrichment of SEC12 mediated by TANGO1 may be achieved by stoichiometric interaction within the TANGO1/cTAGE5/SEC12 complex. One TANGO1 molecule appears to interact with multiple cTAGE5 molecules, which in turn may recruit equivalent amounts of SEC12 (19). Thus, SEC12 may be enriched in several fold molar excess in relation to TANGO1 around ER membrane containing folded PC1.

The cytosolic domains of TANGO1 deleted in our minimal targeting system have also been characterized for their functions during the ER export of PCs. The interaction between the cytosolic PRD domain of TANGO1 and the COPII inner coat subunit SEC23 is proposed to promote the assembly of additional inner coat arrays and stall the recruitment of the outer coat (14, 20). The cytosolic domains of TANGO1 also mediate the formation of TANGO1 rings that are proposed to encircle the necks of budding membranes (45–47). The ring structure is thought to be important for PC secretion and the morphology of the ER and Golgi apparatus.

Sedlin is a small cytosolic protein that interacts with TANGO1 and is required for PC secretion by promoting membrane scission (34). We speculate that the abundance of TANGO1 at the budding neck will recruit more Sedlin to the neck region. Sedlin preferentially binds to SAR1-GTP which stimulates the dissociation of SAR1-GTP from the ER membrane. In light of our discovery that SEC12 extensively covered PC1-rich budding membrane (Fig. 5), membrane scission at the budding neck would be specifically necessary due to the inhibitory effect of SEC12 on vesicle fission (3). We suspect Sedlin is more dispensable for small COPII budding, as SEC12 is excluded from the pre-budding complex.

The COPII outer coat subunit SEC31A is also implicated in the regulation of COPII size and PC1 secretion. SEC31A is a monoubiquitylation substrate of the E3 ubiquitin ligase CUL3, the substrate adaptor KLHL12, and co-factors PEF1 and ALG2 (12, 48). The interaction between SEC31A and KLH12 is important for the monoubiquitylation of SEC31A, enlargement of COPII, and accelerated PC1 secretion (12, 48). Compared to the necessity of TANGO1 during collagen secretion, the KLHL12-SEC31A interaction seems to be more pertinent when timely collagen secretion is required at specific stages during development. As examples, KLHL12 expression was upregulated in embryonic stem cells (12), at specific developmental stages via the UPR transducer BBF2H7 (49), or as a result of CXCL12/CXCR signaling (50). Large COPII vesicles induced by KLHL12 overexpression often exceeded 300nm and appeared less dependent on large cargo, as they can be induced in HEK293T cells that do not express PCs (12). The potential excess of space and sacrifice in selectivity may be beneficial in the interest of speed.

In the present study, we propose a speculative model for the enlargement of COPII-coated vesicles and PC capture coordinated by TANGO1. TANGO1 interacts with HSP47 to detect the presence of folded PC trimers in the ER lumen, and recruits cTAGE5 oligomer which would then enrich SEC12 to the PC1-containing membrane. The concentrated targeting of SEC12 may then promote the formation of larger-than-normal COPII coats by virtue of a persistent recharging of SAR1 on the coated membrane surface. Sorting of PC would thus be ensured by the co-packaging of its adaptor TANGO1 into COPII vesicles, and continued rounds of sorting sustained by recycling the TANGO1-HSP47 complex back to the ERES in COPI vesicles.

## Materials and methods

### Plasmids

Human TANOG1-FLAG in pcDNA3.1 was a gift from Vivek Malhotra lab (CRG-Centre de Regulacio Genomica, Barcelona, Spain). The split GFP plasmids pcDNA3.1 GFP1-10/GFP11 were purchased from AddGene (Cambridge, MA). TANGO1-lumi-HA-GFP1-10 was generated by cloning the luminal domain (1-1142 aa) followed by an HA tag and a linker to the N-terminus of GFP1-10 on pcDNA3.1 GFP1-10. To improve fluorescence intensity of the complemented GFP, we deleted 113-600aa from the luminal domain of TANGO1 with site-directed mutagenesis and used it as TANGO1-lumi-HA-GFP1-10 in this study. Besides improved GFP intensity, results observed using TANGO1-lumi-HA-GFP1-10 with a shortened unstructured domain were indistinguishable from the construct that contained the full-length luminal domain. Human SEC12 in p3xFLAG CMV10 plasmid was a gift from the Kota Saito lab (University of Tokyo, Tokyo, Japan). 3xFLAG-SEC12 was cloned to the N-terminus of GFP11 in pcDNA3.1 GFP11. Site-directed mutagenesis was used to introduce I41A and N43A mutations. HSP47ΔRDEL was cloned from cDNA prepared from U-2OS cells with primers “ATGCGCTCCCTCCTG” and “CATCTTGTCACCCTTAGG” into a pcDNA4 plasmid containing C-terminal strepII tag. The pcDNA4 StrepII plasmid was a gift from the Nevan Krogan lab (UCSF, San Francisco, CA).

### Cell culture, transfection, drug treatment

Human lung fibroblasts IMR-90 and svIMR-90 (IMR-90 immortalized with SV-40) were obtained from Coriell Cell Repositories at the National Institute on Aging, Coriell Institute for Medical Research. Human osteosarcoma Saos-2 and U-2OS and human fibrosarcoma HT-1080 were obtained from ATCC. Because only a low percentage of U-2OS cells express endogenous PC1, we clonally selected U-2OS cells for high endogenous PC1 expression as detected by immunoblotting. The high PC1 expressing clones “wt-C11” and “wt-G5” were used in this work. IMR-90, svIMR-90, Saos-2, U-2OS-wt-C11/G5, and HT-1080 were maintained in DMEM plus 10% FBS (GE Healthcare, Chicago, IL). Dialyzed FBS (Thermo Fisher Scientific, Grand Island, NY) was used to supplement DMEM to make a dialyzed medium. The construction and maintenance of HTPC1.1 and KI6 were described in Jin et al., 2012 and Gorur et al., 2017. HTPC1.1 that stably expresses COL1A1 was constructed from HT-1080, and the doxycycline-inducible KLHL12-3xFLAG stable cell line (KI6) was constructed from HTPC1.1. DNA plasmids were transfected using lipofectamine 2000 according to the manufacturer protocol. Knockdown of ARCN1 was achieved after transfecting siRNA (Qiagen) using lipofectamine RNAiMAX according to the manufacturer protocol. Sequence of siRNA and quantification of knockdown efficiency were described in Sirkis et al., 2017 (26).

### Immunofluorescence, immunoblotting, antibodies

Immunofluorescence (IF) for confocal microscopy and STORM and immunoblotting (IB) analyses were performed as previously described (11,22). For the split GFP experiments, coverslips were mounted in ProLong-Diamond antifade mountant (Thermo Fisher Scientific), which better preserves GFP intensities, overnight before imaging; coverslips for other confocal experiments were mounted in ProLong-Gold antifade mountant (Thermo Fisher Scientific). For immunoblotting detection of TANGO1 in reconstituted COPII vesicles, we used a transfer buffer containing 0.1% SDS. The following antibodies were used: mouse anti-PC1 (clone 42024; QED Biosciences, San Diego, CA; 1:200 for IF); rabbit anti-PC1 (LF-41, 1:5000 for IB) was a gift of Larry Fisher (NIH, Bethesda, MD); rabbit anti-SEC31A (Bethyl Laboratories, Montgomery, TX; for IF, at 1:200 for confocal and 1:2,000 for STORM); StrepMAB Chromeo488 conjugate (IBA Life Sciences, Göttingen, Germany, 1:200 for IF); mouse anti-FLAG (Thermo Fisher Scientific; 1:5,000 for IB); goat anti-FLAG (Novus Biologicals, Littleton, CO; for IF at 1:1,000 for confocal and 1:5,000 for STORM); mouse anti-HSP47 (Enzo Life Sciences Farmingdale, NY; at 1:200 for IF and 1:5,000 for IB); rabbit anti-calnexin (Abcam, Cambridge, U.K., 1:500 for IF, 1:5000 for IB), rabbit anti-TANGO1 (Sigma, St. Louis, MO., 1:200 for IF, 1:1000 for IB); rabbit anti-cTAGE5 (Sigma, 1:2000 for IB); rabbit anti-GFP (Tory Pines, Secaucus, NJ, 1:2000 for IB); goat anti-SEC12 (PREB) (R&D systems, Minneapolis, MN, 1:1000 for IB. Note: this was only used for vesicles purified by density gradient flotation due to a major unrelated cytosolic band); rat anti-SEC12 was purified from hybridoma clone 6B3 provided by the Kota Saito lab (University of Tokyo, Tokyo, Japan) and 1mg/ml aliquots were used 1:1000 for IB and 1:100 for IF.

### Confocal imaging

Confocal imaging was acquired using Zen 2010 Software on an LSM 710 confocal microscope system (ZEISS, Oberkochen, Germany) at CRL Molecular Imaging Center (UC Berkeley, RRID:SCR_012285). The objective used was Plan-Apochromat 63×, 1.4 NA oil DIC M27 (ZEISS). The excitation lines used were 405, 488, 561, and 633nm and collected sequentially with the following spectral range: DAPI (410-481nm), GFP (492-570nm), Alexa568 (570-648nm), and Cy5 (638-755nm) with a pinhole opening of 1a.u. for all channels. Single plane images were acquired at speed 6 for 2048×2048 and averaging of 2 frames. Z stacks of large vesicles were acquired with 0.38µm steps in Z, 2x digital zoom, at speed 8 for 1024×1024 and average of 2.

### STORM imaging

Dye-labeled cell samples were mounted on glass slides with a standard STORM imaging buffer consisting of 5% (w/v) glucose, 100 mM cysteamine, 0.8mg/ml glucose oxidase, and 40µg/ml catalase in a Tris-HCl buffer (pH 7.5) (51, 52). Coverslips were sealed using Cytoseal 60. STORM imaging was performed on a homebuilt setup based on a modified Nikon Eclipse Ti-E inverted fluorescence microscope using a Nikon CFI Plan Apo λ 100x oil immersion objective (NA 1.45). Dye molecules were photoswitched to a dark state and imaged as they individually returned to an emitting state, using either 647 or 560nm lasers (MPB Communications, Quebec, Canada); these lasers were passed through an acousto-optic tunable filter and introduced through an optical fiber into the back focal plane of the microscope and onto the sample at intensities of ∼2 kW cm^-^2^^. A translation stage was used to shift the laser beams towards the edge of the objective so that light reached the sample at incident angles slightly smaller than the critical angle of the glass-water interface. A 405nm laser was used concurrently with either the 647 or 560nm lasers to reactivate fluorophores into the emitting state. The power of the 405nm laser (typical range 0-1 W cm^-^2^^) was adjusted during image acquisition so that at any given instant, only a small, optically resolvable fraction of the fluorophores in the sample were in the emitting state. For 3D STORM imaging, a cylindrical lens was inserted into the imaging path so that images of single molecules were elongated in opposite directions for molecules on the proximal and distal sides of the focal plane (51). The raw STORM data was analyzed according to previously described methods (51, 52). Data was collected at a framerate of 110Hz, for a total of ∼80,000 frames per image.

Three-color imaging was performed on targets labeled by Alexa Fluor 647, CF680, and CF568 via sequential imaging with 647nm and 560nm excitation. With 647nm excitation, a ratiometric detection scheme was employed to concurrently collect the emission of Alexa Fluor 647 and CF680 (53, 54). Emission of these dyes was split into two light paths using a long pass dichroic mirror (T685lpxr; Chroma), each of which were projected onto one-half of an Andor iXon Ultra 897 EM-CCD camera. Dye assignment was performed by localizing and recording the intensity of each single molecule in each channel. Excitation (560 nm) was subsequently used to image CF568 through the reflected light path of the dichroic mirror.

### Vesicle budding reaction

The PC1 budding reaction was performed as described in Gorur et al., 2017 and Yuan et al., 2017. To separate COPII-coated PC1 carriers and regular COPII vesicles, we scaled-up budding reactions 4-fold. In a polycarbonate 11×34mm ultracentrifugation tube (Beckman Coulter), 500µl of 18% (w/v) OptiPrep in B88 (20mM HEPES, pH 7.2, 250mM sorbitol, 150mM KOAc, 5mM Mg(OAc)22) was placed at the bottom of the tube, overlayed with 400µl 7.5% (w/v) OptiPrep (Sigma-Aldrich) in B88, then 350µl of the 7,000xg supernatant of the reaction. The OptiPrep gradient was centrifuged at 250,000xg for 1h at 4°C (Beckman TLS-55) with slow acceleration and deceleration, after which 100µl fractions were collected from the top. Low buoyant density membranes concentrated in the region of 300-400µl from the top (referred to as Fraction 2 in text), and higher buoyant density membranes were collected at position 700-800µl from the top (referred to as Fraction 4 in text). Desired fractions were subjected to flotation: 85 µl of each sedimentation fraction was gently mixed with 50µl of 60% (w/v) OptiPrep until the sample was homogeneous, placed at the bottom of a 7×20 mm tube (Beckman Coulter), and overlaid with 100µl of 18% (w/v) and 10µl of 0% OptiPrep in B88. The OptiPrep gradient was centrifuged at 250,000xg for 1h at 4°C (Beckman TLS-55 with adaptors for 7×20mm tubes) with slow acceleration and deceleration, after which 40µl was collected from the top and mixed with sample buffer for immunoblotting analysis.

### Tryptophan fluorescence assay

The tryptophan fluorescence assay was performed at 37°C in a stirred-cuvette as previously described with slight modifications (3). To HKM buffer (20mM HEPES, pH 7.2, 160mM KOAc, 1mM Mg(OAc)22), an indicated amount of WT, I41A, or N43A SEC12-cyto was added, followed by 2µM SAR1BΔN (the N-terminal amphipathic helix was omitted because it does not affect GTP/GDP binding(4), then 30µM GTP. The fluorescence intensity was followed for 15-20min until the exchange of GDP for GTP equilibrated as detected on a Cary Eclipse Fluorescent Spectrophotometer (Agilent Technologies, Santa Clara, CA) with the following settings: excitation wavelength 297nm; slit size 2.5nm; emission wavelength 340nm, slit size 20nm; high pmt dection (800v); fluorescence reading every 1s for 20min in kinetic mode.

## Supplementary figures with legends

**Figure S1.**
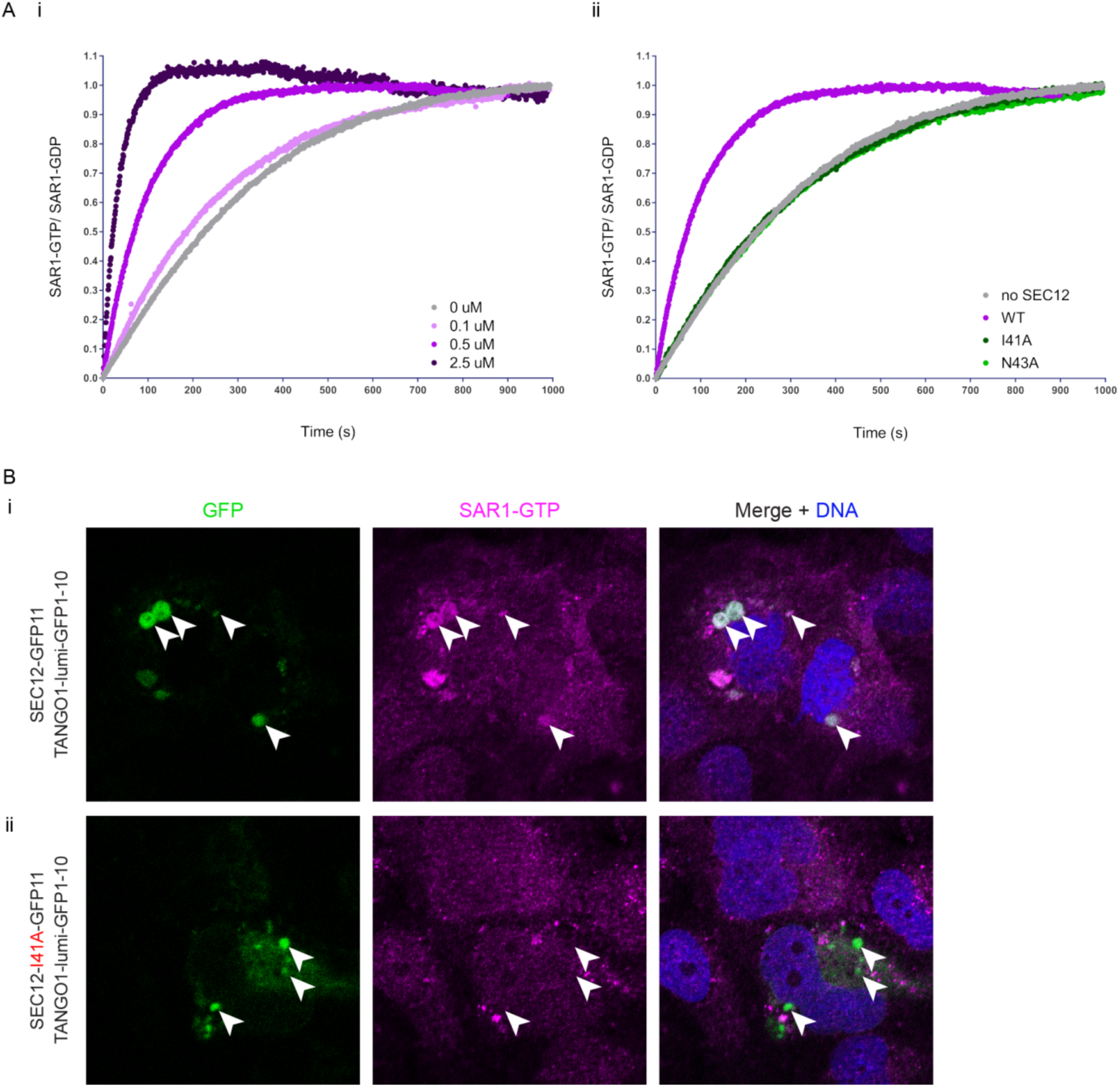
Complemented GFP puncta contain SEC12 with active GEF activity. (A) Mutations in I38 or N40 in Sec12p ablated its GEF activity (3, 35). Thus we mutated the conserved I41 and N43 residues in human SEC12 and tested the GEF activity of wild-type (WT), I41A, and N43A using a tryptophan fluorescence assay. Changes in tryptophan fluorescence intensity were measured with time as an indicator of the nucleotide associated with SAR1 (5). Fraction of SAR1-GTP was calculated by normalizing against the total increase of tryptophan fluorescence after GDP was exchanged to GTP. (i-ii) GDP to GTP exchange occurred naturally (grey) or in the presence of GEF deficient mutants I41A or N43A (green) or was accelerated by WT SEC12 (magenta). Indicated amounts of WT SEC12 were used in (i), and 0.5 µM of WT or mutant SEC12 were used in (ii). (B) Accumulation of SAR1-GTP (magenta) at complemented GFP (green) puncta was observed by confocal microscopy in cells expressing WT SEC12 (i) but not the GEF deficient I41A mutant (ii). U-2OS cells were transfected with both TANGO1-lumi-HA-GFP1-10 and 3xFLAG-SEC12-GFP11 (WT or I41A as indicated), immunofluorescently labeled against SAR1-GTP, and imaged for the complement GFP and labeled SAR1-GTP.

**Figure S2.**
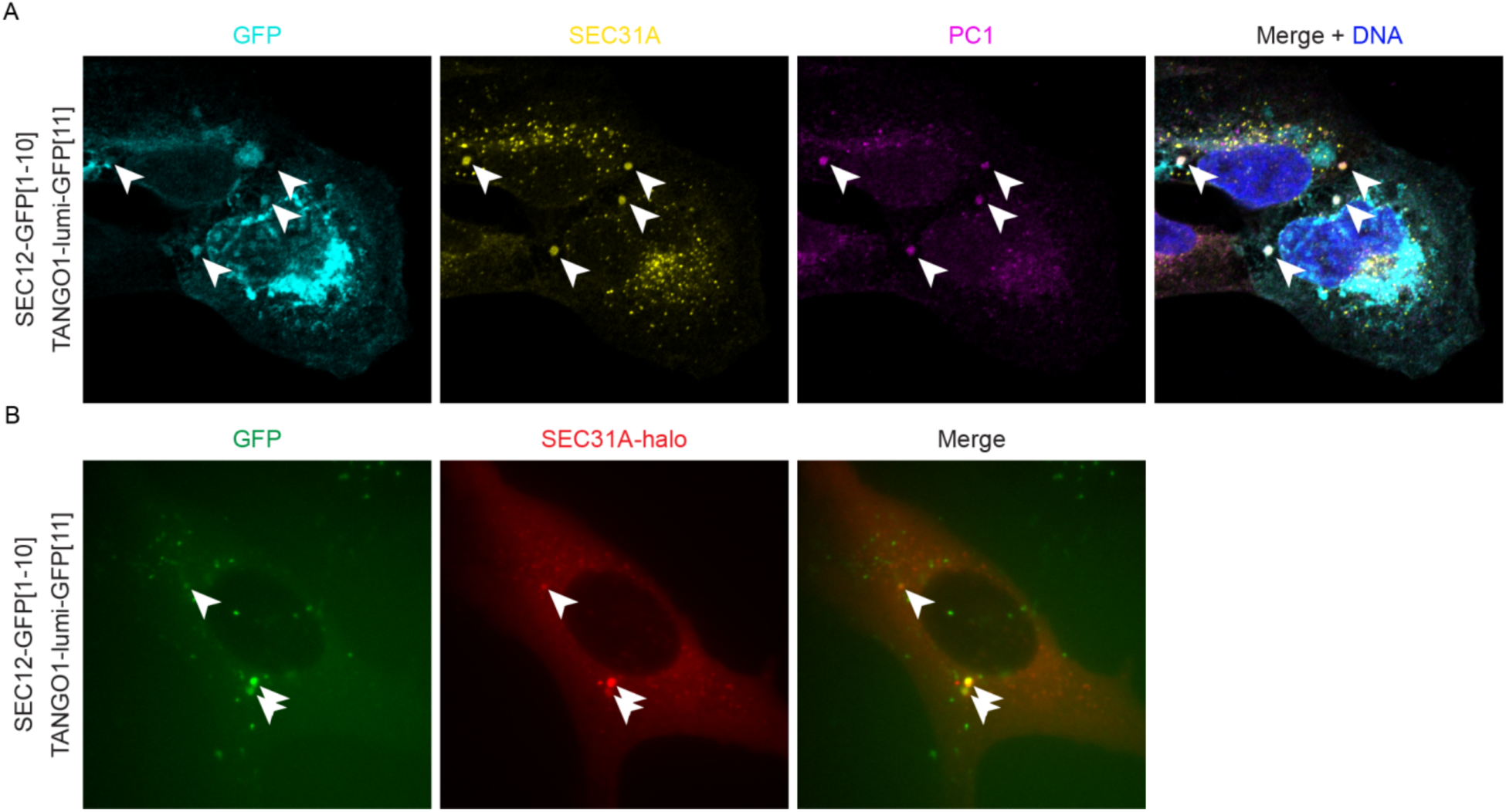
SEC31A colocalizes with large complemented GFP puncta. (A) U-2OS cells were transfected with both TANGO1-lumi-HA-GFP1-10 and 3xFLAG-SEC12-GFP11, then immunofluorescence was visualized by confocal microscopy with PC1 (yellow) and SEC31A (magenta) labels. Occasionally, cells containing large punctate complemented GFP (cyan) signal also show large colocalizing SEC31A and PC1 puncta. (B) U-2OS cells were transfected with SEC31A-halo in addition to TANGO1-lumi-HA-GFP1-10 and 3xFLAG-SEC12-GFP11. Live cells were imaged on an epifluorescence microscope after SEC31A-halo was labeled with the halo ligand TMR (red). Large puncta that contained complement GFP (green) and SEC31A (red) were observed at very low frequency.

### Acknowledgment

We thank the staff at UC Berkeley shared facilities including Alison Killilea (UC Berkeley Cell Culture Facility) and Holly Aaron (UC Berkeley CRL Molecular Imaging Center). Confocal microscopy experiments reported in this publication was performed at CRL Molecular Imaging Center, supported in part by the Gordon and Betty Moore Foundation. We thank Jeremy Thorner for providing the fluorometer and Shawn Shirazi for 3D printing that was essential for the setup. We thank Vivek Malhotra for advice and discussion during his sabbatical at the Schekman lab. We also thank past and present members of the Schekman Lab, in particular, Amita Gorur, Dan Sirkis, Liang Ge, and Johannes Freitag. R.S. is supported as an Investigator of the Howard Hughes Medical Institute and the UC Berkeley Miller Institute of Science. L.Y. was supported in part by the Tang family fellowship. S.K. and K.X. acknowledge support from NSF under CHE-1554717, the Pew Biomedical Scholars Award, and the Chan Zuckerberg Biohub. The authors declare no competing financial interests.

## Author contribution

L.Y., K.X., R.S., designed research; L.Y., S.J.K, J.H., performed research, contributed new reagents or analytic tools; L.Y., S.J.K, K.X., R.S. analyzed data, and wrote the paper.

